# The cell cycle state defines TACC3 as a regulator gene in glioblastoma

**DOI:** 10.1101/2020.10.20.346643

**Authors:** Holly Briggs, Euan S. Polson, Bronwyn K. Irving, Alexandre Zougman, Ryan K. Mathew, Deena M.A. Gendoo, Heiko Wurdak

## Abstract

Overexpression and mitosis-promoting roles of Transforming acidic coiled-coil containing protein 3 (TACC3) are well-established in many cancers, including glioblastoma (GBM). However, the effector gene networks downstream of TACC3 remain poorly defined, partly due to an incomplete understanding of TACC3 cell lineage specificity and its dynamic role during the cell cycle. Here, we use a patient-derived GBM model to report that TACC3 predominantly resides in the GBM cell cytoplasm, while engaging in gene regulation temporally as defined by the cell cycle state. TACC3 loss-of-function, cell cycle stage-specific transcriptomics, and unsupervised self-organizing feature maps revealed pathways (including Hedgehog signalling) and individual genes (including HOTAIR) that exhibited anticorrelated expression phenotypes across interphase and mitosis. Furthermore, this approach identified a set of 22 TACC3-dependent transcripts in publicly-available clinical databases that predicted poor overall and progression-free survival in 162 GBM and 514 low-grade glioma patient samples. These findings uncover TACC3-dependent genes as a function of TACC3 cell cycle oscillation, which is important for TACC3-targeting strategies, and for predicting poor outcomes in brain cancer patients.

## Introduction

Elevated expression of Transforming Acidic Coiled-Coil Containing Protein 3 (TACC3) has been associated with adverse patient outcomes in a wide range of different cancers including brain, breast, and lung (1). Following the first functional characterization of *TACC3 in vivo* (2), an ever-increasing number of studies implicate the expression of TACC3 (3–8) and/or TACC3 fusion proteins (e.g., TACC3-FGFR3; (9)) in tumor growth. Consistently, TACC3 has been reported to regulate the expression of transcription factors that promote cell proliferation, including E2F Transcription Factor 1 (E2F1; (10)) and MYC Proto-Oncogene, BHLH Transcription Factor (MYC) (11). However, despite the evolving notion of a TACC3-dependent axis of oncogenic gene expression, many molecular pathways downstream of elevated TACC3 are not yet understood. TACC3 expression has been described as both cell lineage-specific and dependent on the cell cycle (2), and mechanistic TACC3 roles at the centrosome and mitotic spindle (12–14), and the cytoplasm (during interphase (15)) have emerged. To make matters even more complex, other non-mitotic roles of TACC3 have been described, including neural differentiation (16), cytoplasmic transcription factor sequestration (17), signaling (8,18,19), and ciliogenesis (5). This raises questions as to whether TACC3 multifunctionality is linked with an equally dynamic downstream gene expression network in tumor cells, and whether the cell cycle state matters for TACC3 gene-regulatory activities. As transcriptomic analysis of TACC3 temporal effects and cell-state specific data are very limited, we sought to address this question utilizing a TACC3 RNA interference (RNAi) loss-of-function approach in a patient-derived cell model of the most frequent and aggressive primary brain cancer, glioblastoma (GBM) (20). We determined the TACC3 localization and expression patterns across the cell cycle and carried out cell cycle phase transcriptomics after TACC3 knockdown, followed by unsupervised data clustering through self-organizing feature maps (SOM) (21). The results uncover TACC3 as a conditional regulator gene across cell cycle phases, bringing to light the pathways and prognostic genes that are under periodic TACC3 control in the GBM cellular context.

## Results

### TACC3 localization is predominantly cytoplasmic in cancer cells

As an initial step, we examined TACC3 cytoplasmic and nuclear localization patterns using publicly available immunohistology data (22) (Fig. 1A, Supplementary file 1: Table S1). Analysis of histological tissue sections from four different cancer types (brain, breast, lung, lymphoma) showed that TACC3 was predominantly localized in the cytoplasm of the malignant cells (>50% of at least 1000 cells analyzed per section, Fig. 1A). Next, we assessed TACC3 localization and expression using our patient-derived primary GBM cell model GBM1 (20). GBM1 cells exhibit a mixed classical/mesenchymal glioblastoma subtype profile and a tumor stemness/proliferation phenotype that is abrogated upon TACC3 downregulation (20). Consistent with a high cytoplasmic TACC3 content in cancer tissues, GBM1 cells showed a marked increase in cytoplasmic TACC3 that was concomitant with reduced nuclear protein when compared with the HeLa cell line, which has been widely used for TACC3 characterization (a literature search for ‘TACC3, HeLa’ retrieved >30 PubMed entries) (Fig. 1B). Moreover, TACC3 mRNA expression was markedly elevated in GBM1 cells (>4-fold, Fig. 1C). TACC3 interaction proteomics in GBM1 and HeLa cells indicated wide-spread cytoplasmic and nuclear TACC3 physical association networks (Fig. 1D). In addition, the expected cell type specificity of the TACC3 interactome was observed in GBM1 cells as exemplified by the presence of (cytoplasmic) intermediate filament proteins Nestin (NES) and Vimentin (VIM), which have both been linked to brain tumor progression (24,25). Taken together, these data identify GBM1 as a suitable model for studying malignant TACC3 overexpression.

**Figure 1.**
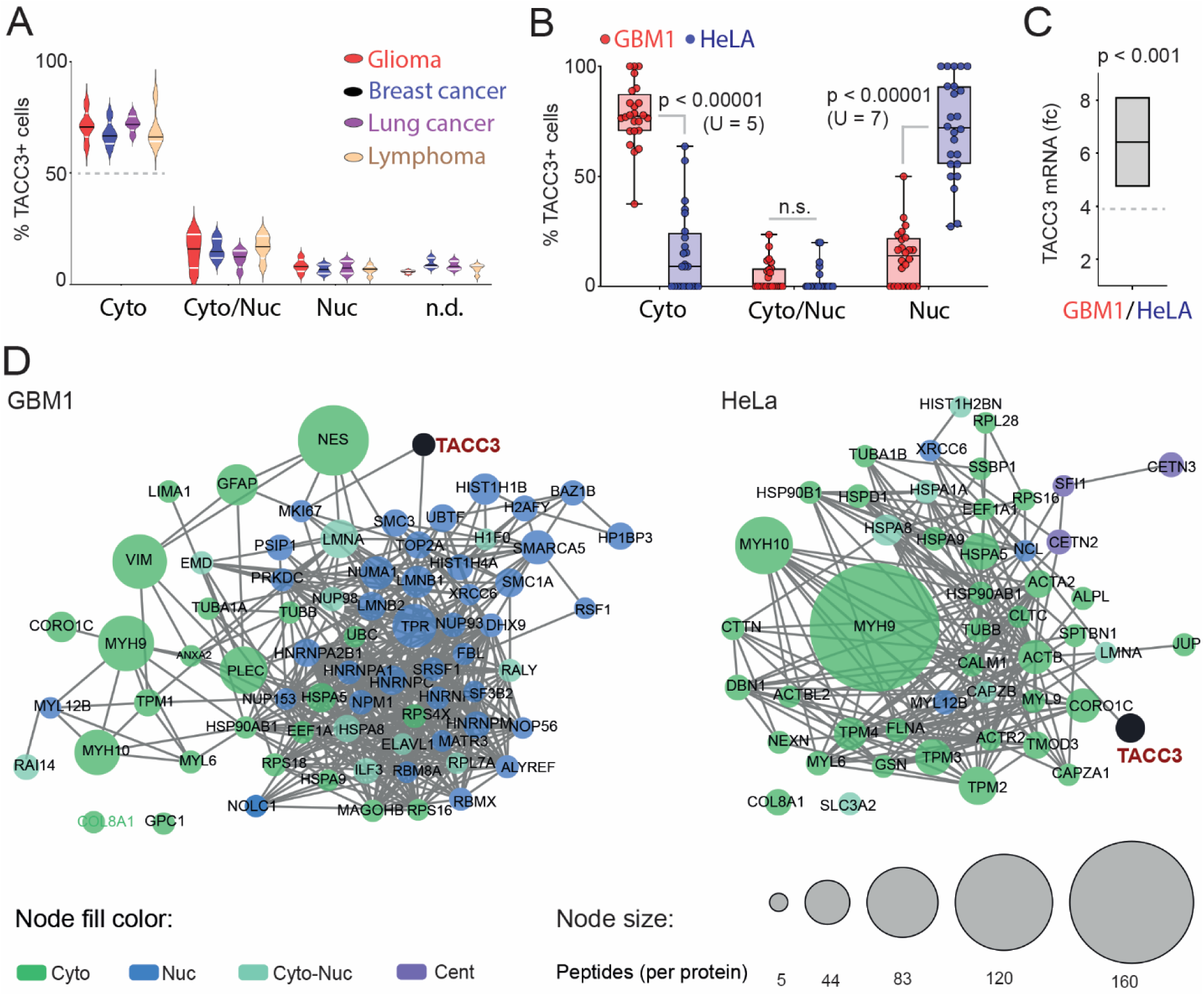
TACC3 localization is predominantly cytoplasmic in cancer cells. **A)** Percentage of cancer cells in tissue sections of the indicated cancer types (n = 5; http://proteinatlas.org) staining positive for TACC3 exclusively in the cytoplasm (Cyto), both cytoplasm and nucleus (Cyto/Nuc), and only nucleus (Nuc); n.d.: localization not determined. **B)** Box plots (minimum to maximum) of distinct TACC3 cytoplasmic (Cyto), cytoplasmic/nuclear (Cyto/Nuc), and nuclear (Nuc) staining (percentages) in GBM1 and HeLa cells (red and blue dots). **C)** Intercleaved (low-high) plot of relative TACC3 mRNA expression in GBM1 versus HeLa cells; n = 3, fc: fold-change.**D)** Cytoscape (19) visualization of the TACC3-specific immunoprecipitated proteins identified by proteomics; Node size: total peptides detected per indicated protein. Network edges indicate protein relationships as designated in (23). Network nodes are colored by the subcellular location for each detected TACC3-associated protein: cytoplasm (Cyto), nuclear (Nuc), centrosomal (Cent), and cytoplasmic/nuclear (Cyto/Nuc).

### TACC3 expression and downstream gene-regulatory activities are dependent on the cell cycle stage

Stable expression of the FUCCI indicator (26) in GBM1 cells (GBM1_FUCCI) enabled monitoring of TACC3 expression and localization across the early G1 (eG1), G1, S, and G2/M cell cycle phases. Cytoplasmic TACC3 showed an oscillatory pattern, reaching peak levels during the G2/M phase, whereas nuclear TACC3 was exclusively observed in G2/M fractions of GBM1_FUCCI cells after sorting (Fig. 2A). Expression of both TACC3 mRNA and protein oscillated with cell cycle progression, reaching lowest levels during G1, and highest levels during G2/M (Fig. 2B). Next, we employed lentiviral TACC3 RNAi using two different small hairpin RNA knockdowns (Kd1, Kd2) that resulted in a significant reduction in TACC3 mRNA and protein levels after 48 hours in GBM1_FUCCI cells (Fig. 2C, Supplementary file 1: Fig. S1). We transcriptionally profiled each cell cycle stage relative to the negative control. Potential RNAi off-target effects were minimized by focusing our downstream analysis on differentially expressed genes that showed phenotypic redundancy (high probe value correlation) between Kd1 and Kd2 (Supplementary file 2). Principal component analysis (PCA) revealed TACC3-dependent gene expression as a function of cell cycle stage (Fig. 2D), and subsequent single-sample gene set enrichment analysis (ssGSEA) confirmed cell cycle-dependent enrichment of TACC3 relevant cell cycle-, cell morphology-, and cancer-associated pathways (Supplementary file 3, Fig. 2E). Pertinent pathways include cytoskeleton organization and G2/M checkpoint control pathways (expected to positively enrich toward G2/M), and the Aurora kinase A (AURKA) pathway (expected to show enrichment correlating with TACC3 expression over the cell cycle, with TACC3 being a known phosphorylation target of AURKA (27)).

**Figure 2.**
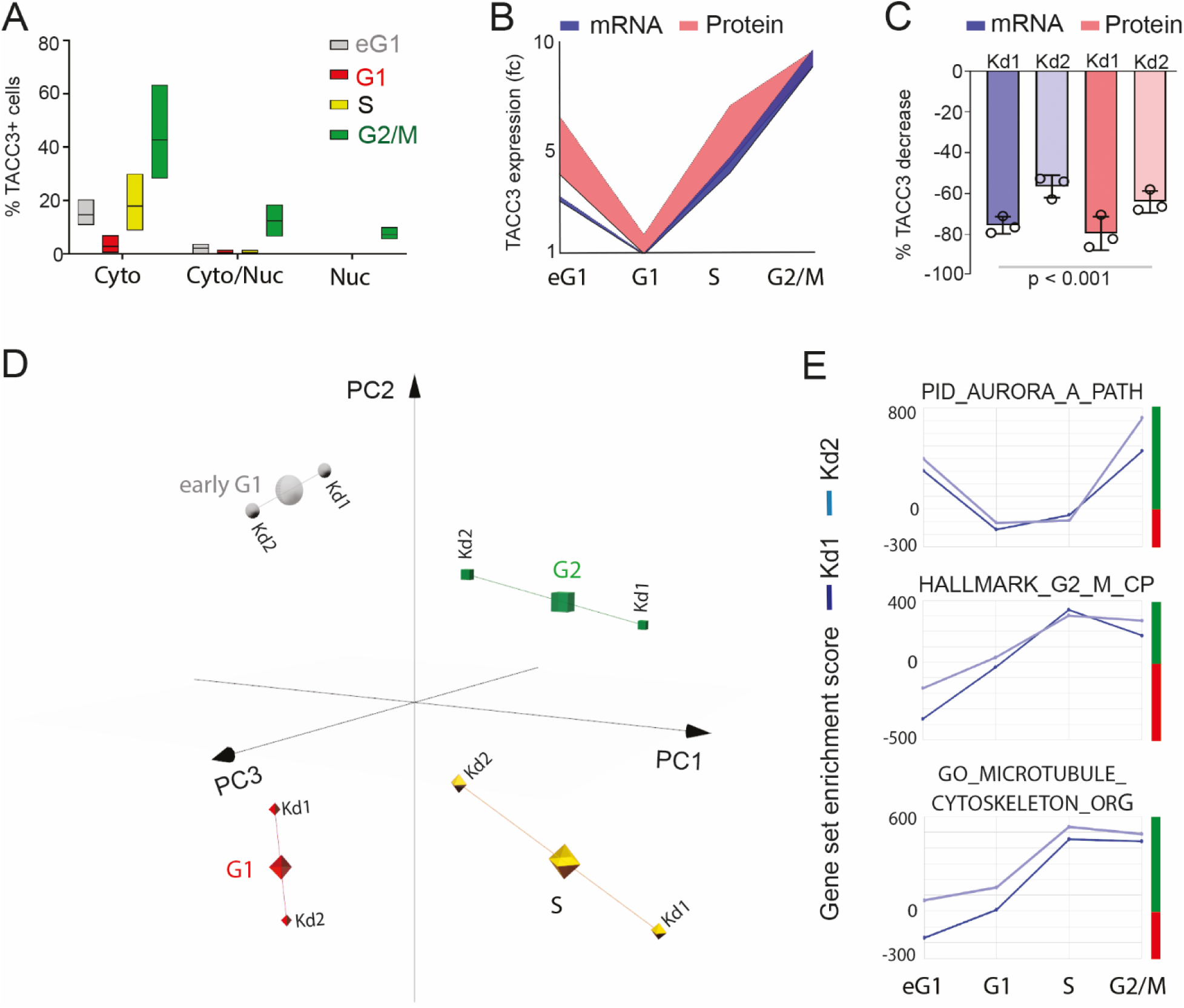
TACC3 expression and downstream gene-regulatory activities are dependent on the cell cycle stage. **A)** Interleaved (low-high) plots of cytoplasmic (Cyto), cytoplasmic and nuclear (Cyto/Nuc), and nuclear (Nuc) TACC3 staining (percentages) across the indicated cell cycle phases in GBM1-FUCCI cells. **B)** Area fill plot of mRNA and protein levels across the indicated GBM1_FUCCI cell cycle phases. **C)** Change in TACC3 mRNA and protein levels post TACC3 knockdowns (Kd1, Kd2) relative to the control in GBM1-FUCCI cells; percent decrease; n = 3, p-value: one-way ANOVA. **D)** PCA analysis (showing first three principal components) of GBM1_FUCCI cell cycle transcriptional profiling upon TACC3 knockdown (Kd1, Kd2); cluster centroids indicated by larger symbol. **E)** ssGSEA analysis of GBM1_FUCCI cell cycle transcriptional profiling showing positive (green bar) and negative (red bar) enrichment scores for the indicated pathway examples.

### Unsupervised clustering places TACC3 upstream of a gene regulatory network that dynamically changes with cell cycle progression

Based on ssGSEA analysis of TACC3-dependent gene expression, we sought to visualize the biological processes that had been most dramatically affected by TACC3 loss-of-function in the GBM1_FUCCI model. Using the ssGSEA-derived enrichment scores (ES) from 142 pathways, across the eG1, S and G2/M stages, we used an unbiased data clustering approach through SOMs (21). Categorization of the ssGSEA pathways (based on their respective ES per cell cycle stage) was subsequently assigned to a map grid composed of 20 nodes and 10 clusters (Path_SOM, Fig. S2A), with individual clusters indicating the most pronounced ES dynamics as a function of cell cycle transitions.

Our Path_SOM highlighted seven nodes that formed individual clusters (Fig. 3A, Fig. S2B). These clusters (c3, c9, c5, c16, c10, c4, c11) harbored 38 pathways that were characterized by their TACC3-dependent cell cycle phenotypes, including: eG1 downregulation (c3), G2/M upregulation (c9, c5), and G2/M downregulation (c16) as compared with all other cell cycle phases. In addition, pathways were categorized by their general downregulation (c10, c4) and upregulation phenotypes (c11) (Fig. S2B). Consistent with the observation that TACC3 levels escalate during the second half of the cell cycle (Fig. 2A), Path_SOM highlighted 19 pathways (including cell division and mitotic checkpoint gene sets) sustaining moderately-positive ES in the presence of TACC3 RNAi during S and G2/M, but not eG1/G1 phases (clusters c3, c9, c5; Fig. 3A and B). In contrast, 7 pathways were upregulated by TACC3 loss-of-function during the bulk of interphase (eG1/G1/S) compared with the G2/M stage (c16; Fig. 3A and B). This cluster indicates a temporal regulatory role for TACC3 in adherens junction and Hedgehog signaling pathway suppression. Path_SOM also highlighted 4 pathways (including TGFβ signaling) that were persistently upregulated across the GBM1_FUCCI cell cycle by TACC3 RNAi (c11, Fig. 3A and B). In contrast, 8 pathways (including DNA/RNA synthesis-fueling pyrimidine and purine metabolism pathways) were downregulated (c10, c4; Fig. 3A and B). Together, these results demonstrate that TACC3 dynamically regulates the expression of both interphase and mitosis-promoting gene networks in a cell cycle-dependent fashion in GBM1_FUCCI cells.

**Figure 3.**
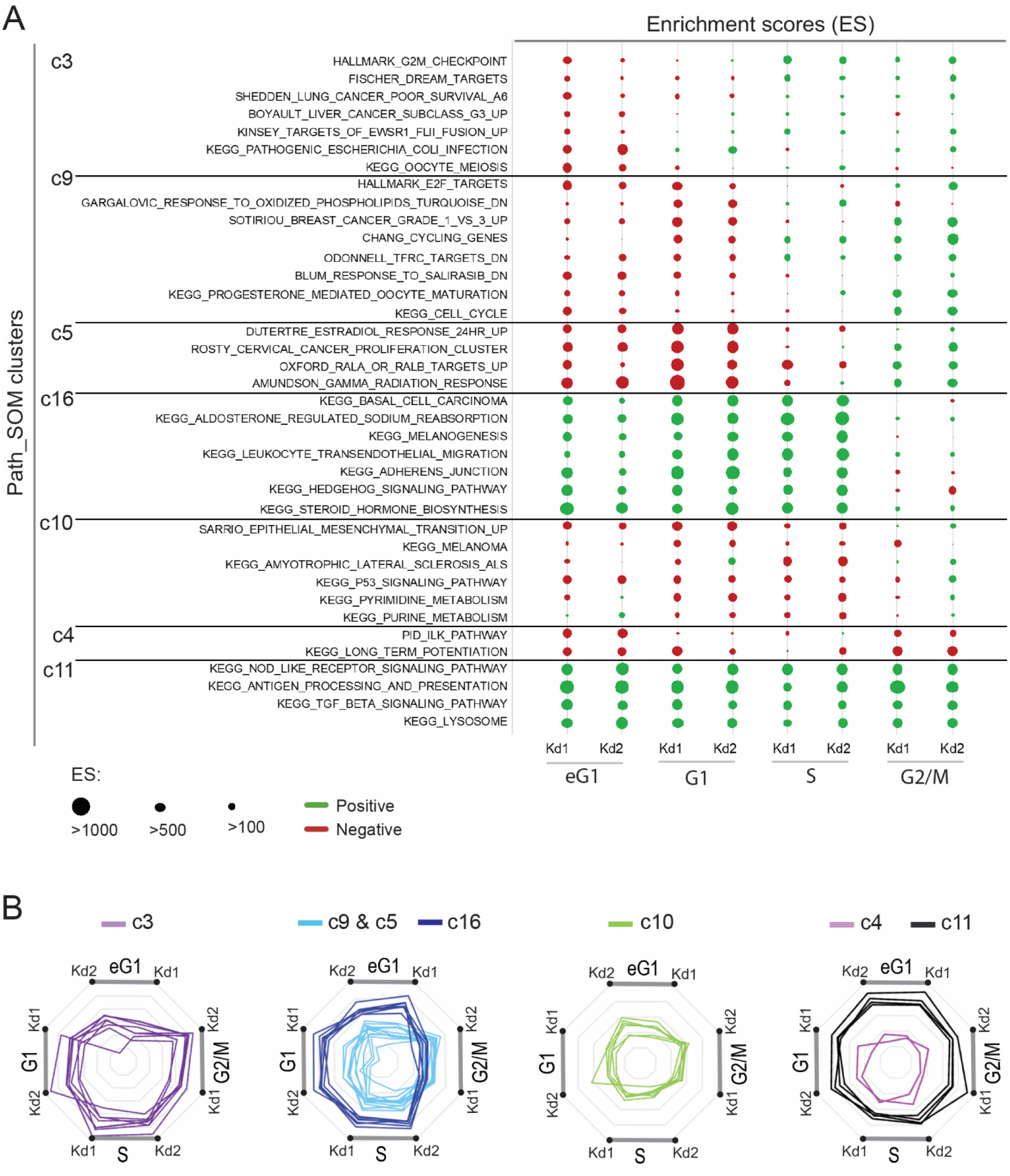
Unsupervised clustering places TACC3 upstream of a gene regulatory network that dynamically changes with cell cycle progression. **A)** Enrichment scores (ES) of TACC3-dependent pathways within individual Path_SOM clusters (see also Figure S3). Pathway ES are shown for both TACC3 knockdowns (Kd1, Kd2) over the depicted cell cycle stages. **B)** Radar plots for pathways listed in (a) based on ES (Additional file Each of the indicated cell cycle phases and TACC3 knockdowns (Kd1, Kd2) are shown; eG1: early G1.

### TACC3 is upstream of genes associated with poor GBM and LGG survival

Encouraged by the Path_SOM results, we used a similar approach (Gene_SOM) to identify a brain tumor-specific signature of TACC3 cell cycle-dependent gene expression (TACC3-CCDGE) that might be useful for predicting overall survival. Unsupervised mapping of normalized TACC3 Kd1 and Kd2 expression values over the GBM1_FUCCI cell cycle identified 238 genes that were represented within 7 individual clusters (Supplementary file S1: Table S2 and Fig. S3A). We further filtered for the most suitable TACC3-CCDGE candidate genes by assessing whether their elevated expression negatively correlated with GBM patient survival (Logrank p-value < 0.05 as determined via (28)). Twenty-two genes in 6 of the 7 clusters fulfilled these criteria (Fig. 4A and Supplementary file S1: Table S3). Among those, 8 genes were associated with Logrank p-values below 0.015 including Lysyl oxidase (LOX), Alkaline phosphatase, Biomineralization Associated (ALPL), Prolyl-4-hydroxylase subunit 2 (P4HA2), and HOX Transcript Antisense RNA (HOTAIR), which have all been previously implicated in which have all been previously implicated in brain tumor biology (29–32). HOTAIR stands out as an oncogenic long non-coding RNA (lncRNA) in different cancers (33), and a promising diagnostic and prognostic biomarker for GBM based on its detectability in the blood of GBM patients (34). Intriguingly, our Gene_SOM clustering revealed HOTAIR as a factor that is temporally regulated by TACC3, as observed by the G2/M-specific HOTAIR downregulation after TACC3 knockdown compared with eG1/G1/S phases (c21, Fig. 4A).

**Figure 4.**
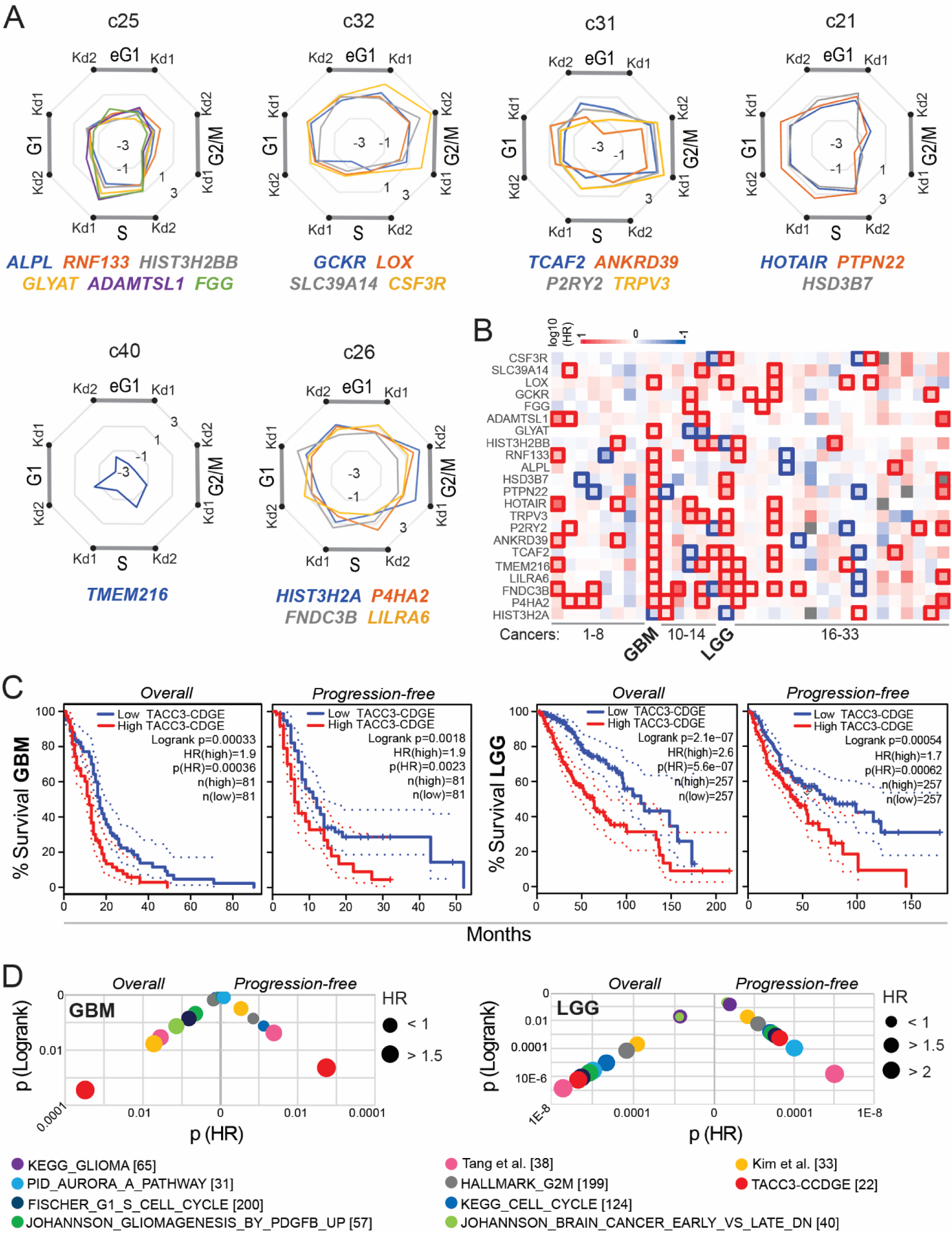
TACC3 is required for gene expression associated with poor GBM and LGG survival. **A)** Radar plots of six individual clusters (c25, c32, c31, c21, c40, c26) identified by unsupervised SOM analysis (Gene_SOM; see Supplementary Figure 4). Indicated genes for each cluster are genes differentially regulated by TACC3 knockdowns (Kd1, Kd2) across the indicated cell cycle phases in GBM1_FUCCi cells, and which are negatively correlated with survival (p < 0.05). Plotted data are fold changes (−3<fc<3) of respective gene expression values (Additional file 2). **B)** Survival map comparing 33 cancer types (Supplementary file 1: Table S4) based on log10 hazard ratios (HR) for the 22 genes forming the TACC3-CCDGE signature. **C)** Kaplan Meier plots showing relationships between the TACC3-CCDGE signature and percent survival for GBM and LGG cohorts. **D)** Bubble plot demonstrating Logrank and hazard ratio (HR) survival across GBM and LGG cohorts, using TACC3-CCDGE and other selected signatures (Supplementary file 1: Table S5). The number of genes per signature is shown in brackets.

Analogous to pathways (as shown in Fig. 3), the expression of the 22 TACC3-CCDGE candidate genes was characterized by cell cycle phenotypes comprising: S phase upregulation (c25), S phase downregulation (c32), G2/M upregulation (c31), G2/M downregulation (c21), general downregulation (c40), and general upregulation (c26) (Supplementary file 1: Fig. S3b). Combined as a 22-gene signature, increased expression of the TACC3-CCDGE set was accompanied with the highest risk of shortened survival (p < 0.05) in the GBM and LGG (brain tumor) contexts as compared to a panel of 31 non-neural (e.g., breast, lung, and colon) cancer types available (35) (Fig. 4B, Supplementary file 1: Table S4). Furthermore, TACC3-CCDGE predicted poor survival in 162 GBM and 514 LGG publicly-available brain tumor specimens with Logrank p-values below 0.0004 for GBM, and 0.002 for LGG, for the assessed ‘Overall survival’ and ‘Progression-free’ categories (as determined via (35)) (Fig. 4C). Compared with 9 selected cell cycle-, TACC3- and brain tumor-relevant gene sets (including those inspired by the ‘Tang et al’, and ‘Kim et al.’ signatures (36,37) and supplied under (35); Supplementary file 1: Table S5), TACC3-CCDGE ranked first with regards to predicting poor survival in GBM, and was among the top three risk predictors in the LGG data set (Fig. 4D). In summary, these results establish TACC3-CCDGE as a gene set that may aid in predicting a poor prognosis in brain tumor patient cohorts.

## Discussion

TACC3 has a well-acknowledged role as centrosomal adapter protein (13) and as a mitotic spindle assembly factor (12,14); however, our understanding of TACC3 during interphase remains limited. This study began with the observation of an overt cytoplasmic TACC3 localization in GBM cell culture and in brain, breast, and lung cancer tissue sections. The dominant presence of TACC3 in the cytoplasm is in agreement with a recent study reporting TACC3 as a regulator of cytoplasmic microtubule plus-end growth, stability, and cargo sorting in interphase cells (15). Moreover, cytoplasmic TACC3 has been shown to sequester transcriptional co-factors away from the nucleus, which can direct progenitor cell fate in a cell type and context-dependent fashion (16,17). Here, the TACC3 physical interactome observed in the GBM1 model suggests that TACC3 may influence GBM-relevant gene expression through diverse interactions. These may include proliferation-associated factors (e.g., MKI67), DNA topology-controlling enzymes (e.g., TOP2A), Lamins (e.g., LMNA) and nuclear pore-associated proteins (e.g., TPR). Further research, independent of the current study, will be required to fully elucidate the spatiotemporal dynamics of both cytoplasmic and nuclear TACC3-protein interactions in diverse cell types. Our findings demonstrate the importance of cell cycle progression for TACC3 expression dynamics in a GBM cellular context. Based on the observed oscillation of TACC3 mRNA and protein expression levels in GBM1_FUCCI cells, we used (TACC3-dependent) gene expression as a readout. Cell cycle state-dependent transcriptional profiling via the GBM1_FUCCI model, and unbiased data clustering through SOMs, enabled us to pinpoint the molecular networks and individual genes downstream of TACC3. Notably, most genes/pathways were periodically dependent on TACC3 during cell cycle progression. One example was Hedgehog signaling as indicated by the positive gene enrichment during early G1, G1, and S (versus the G2/M) phase after TACC3 knockdown. This anticorrelated expression pattern indicates suppression of the Hedgehog pathway during most of interphase and is consistent with the reported TACC3-dependent prevention of premature cell cycle exit in neurogenesis (9), and with TACC3 promoting prostate cancer cell proliferation while restraining the formation of primary cilia (5) that are essential for mammalian Hedgehog signaling (38)).

TACC3 can regulate oncogenic gene expression upstream of transcription factors that promote malignant proliferation such as E2F1 (10) and MYC (11). Here, TACC3 loss-of-function revealed a yet unidentified link between TACC3 and *HOTAIR* expression (indicated by the TACC3 knockdown-induced G2/M downregulation phenotype). Notably, *HOTAIR* is a direct target of MYC in gallbladder cells (39), and a body of literature described HOTAIR as a potential prognostic biomarker and therapeutic target in GBM and other types of cancer (31,33,34,40). One established function of the HOTAIR lncRNA is epigenetic gene regulation via its interaction with Polycomb Repressive Complex 2 (33), hence raising the possibility of a TACC3-HOTAIR gene regulatory axis that is governed by cell cycle dynamics. For example, HOTAIR can inhibit mineralization in osteosarcoma cells by epigenetically repressing *ALPL*, a gene under temporal TACC3 control in GBM1-FUCCI cells (indicated by a distinct S-Phase upregulation phenotype in the absence of TACC3). Both *ALPL* and *HOTAIR* were previously implicated in brain tumor progression (30,31,34,40), and among 22 genes identified by unbiased Gene_SOM clustering (indicating the most differential individual gene expression changes over cell cycle progression), In addition, the expression of these genes significantly correlated with poor GBM survival. Several studies utilized cell cycle-independent transcriptional profiling data to identify gene signatures of potential prognostic value in GBM (36,37). Our results identify the 22-gene-comprising (cell cycle-dependent) TACC3-CCDGE probe set that predicts GBM and LGG ‘overall’ and ‘recurrence-free’ survival compared with 31 other tumor types (35). In this setting, TACC3-CDGE had predictive power compared to several cell cycle- and glioma-focused gene sets (e.g., HALLMARK HALLMARK_G2M_CHECKPOINT and KEGG_GLIOMA). False negative results were mitigated by two probe sets, which were based on previously published GBM survival-indicating signatures (‘Tang et al., Kim et al.’), and these also predicted brain cancer survival in our analysis setting (e.g., ‘Tang et al.’ was associated with the highest significance levels predicting LGG survival). It is worth noting that TACC3-CDGE identification is linked to the herein described cellular context, and data analysis and filtering criteria may vary in different cellular brain tumor and cancer contexts. The potential heterogeneity of TACC3 cell cycle-dependent gene expression networks, which may include the function of TACC3 fusion protein(s), remains to be addressed.

In conclusion, our findings demonstrate the importance of the cell cycle state for genes whose expression requires TACC3 function. Importantly, they place TACC3 upstream of genes with potential prognostic value in brain cancer. Our analysis bypasses limitations of cell cycle-independent profiling, to shed new light on TACC3-dependent effectors and pathways with anticorrelated expression patterns that are linked to individual cell cycle stages. We propose that TACC3-targeting strategies should consider the cell cycle and our TACC3-dependent signature may serve as a readout for further validation.

## Materials and Methods

### Cell culture

Adherent culture of patient-derived GBM cells has been previously described (20). Briefly, GBM1 and GBM1_FUCCI cells were cultured as adherently-growing cells in Neurobasal medium (Gibco) supplemented with human recombinant bFGF, EGF (Gibco, 40 ng/mL each), 0.5x B27, and 0.5x N2 (both Gibco) at 37^°^C with 5% CO2 on poly-L-ornithine/laminin-coated (5 μg/mL each) plastic cell culture flasks or dishes. HeLa cells were obtained American Type Culture Collection (ATCC CCL-2) and cultured in DMEM/F12 supplemented with 10% (v/v) fetal bovine serum.

### TACC3 expression analyses

#### Immunocytochemistry

after fixation with 4% (w/v) paraformaldehyde (10 minutes, RT), cells were probed for TACC3 antibody (Cell Signaling, #80659S, 1:500). Non-specific antibody binding was reduced by incubating fixed cells in PBS ‘blocking buffer containing 10% (v/v) FBS and 0.03% (v/v) Triton X-100 at RT for 1 hour. Primary antibody was used in ‘blocking buffer’ at 4^°^C overnight. Secondary antibody: Cy3 or Cy5-conjugated (Jackson ImmunoResearch; 1:400). Nuclei were stained using DAPI (Sigma; 1 μg/mL). Images were acquired using an EVOS digital inverted fluorescence microscope (life technologies) or a Nikon A1R confocal microscope.

#### qRT-PCR analysis

total RNA was extracted using the RNeasy mini kit (Qiagen) according to the manufacturer’s instructions. Copy DNA was synthesized using the SuperScript II first-strand-synthesis method with oligo(dT)s (Invitrogen) and analyzed using TaqMan gene expression assays (TaqMan Assay ID Hs00170751_m1). Data were internally normalized to Glyceraldehyde-3-phosphate dehydrogenase (GAPDH).

#### Western blotting

complete cellular extracts (~106 cells) were lysed in CellLyticMT (Sigma; C3228) plus protease/phosphatase inhibitor cocktail (Thermo Scientific; 78440) following the manufacturer’s recommendations. To achieve nuclear/cytoplasmic separation of total cellular protein, the CelLytic NuCLEAR Extraction Kit (Sigma; NXTRACT-1KT) was used. Lysate concentrations were determined using the BCA protein assay (Pierce; 23225). Protein (between 15-30 μg) was loaded onto Mini-Protean TGXTM precast gels (10%; Biorad), and transferred onto a nitrocellulose membrane (0.45 μm, Biorad). Membranes were exposed to the following antibodies: mouse anti-TACC3 (Santa Cruz; sc-48368; 1:500), mouse anti-Actin beta (Sigma; A5441; 1:20,000).

#### FACS

GBM1 cells stably expressing FUCCI were washed with PBS and detached from the plate using Accutase (STEMCELL Technologies #07922) for ~10 min at 37ºC. Cells were washed twice with PBS and subjected to cell sorting using parameters that were previously described (7). A FACS Aria (BD Biosciences) cell sorter was used, and non-transduced GBM1 cells served as a gating control.

#### Tissue sections

digital images of TACC3 protein expression within five sections per cancer type were obtained from the Human Protein Atlas database (https://www.proteinatlas.org/ENSG00000013810-TACC3/pathology) (22). Images were analyzed using ImageJ software and total nuclei as well as TACC3 immunopositivity and (cytoplasmic/nuclear) localization were determined by averaging the quantifications from two different investigators.

### TACC3 RNAi

Generation of lentiviral particles was carried out according to the manufacturer’s specifications and as previously described (20). After 16 hours, medium containing residual virus was washed out and replaced with fresh media containing 2 μg/mL puromycin. Protein and mRNA knockdown (Kd) efficiencies were determined by Western blotting and qRT-PCR at 48 hours post viral transduction. The two different lentiviral TACC3 small hairpin (sh) RNA and non-targeting constructs were:

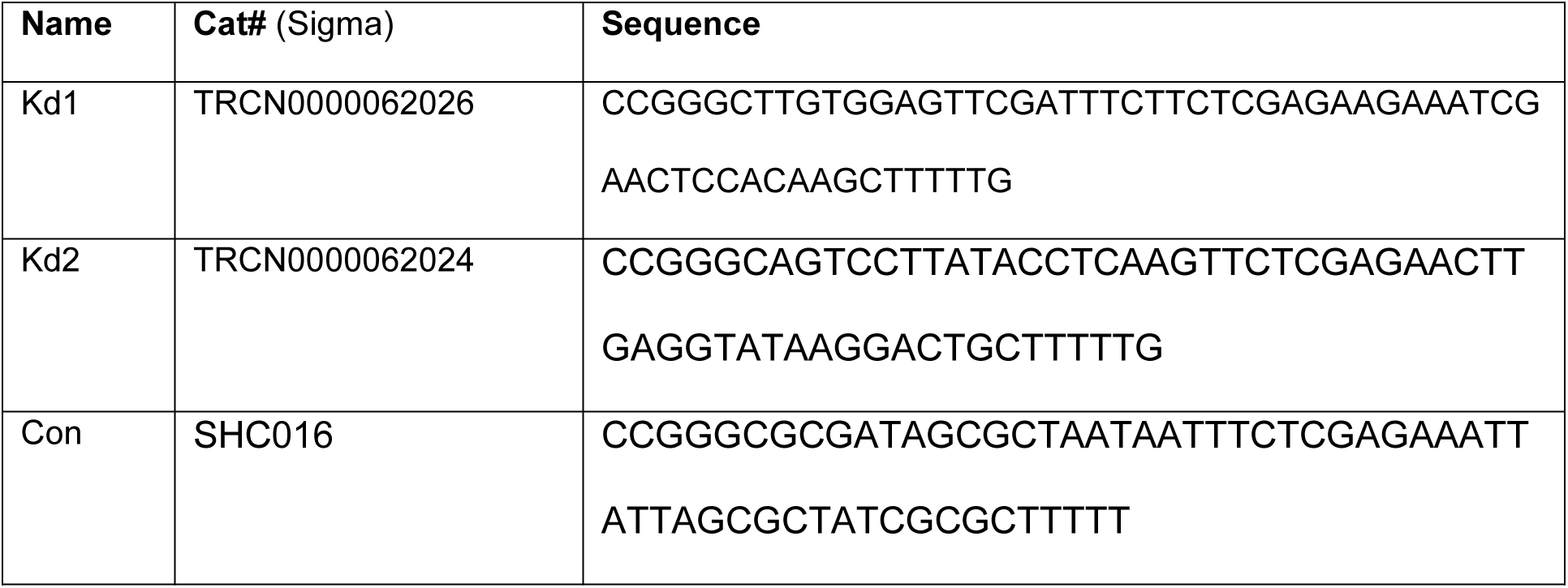

### Proteomic analysis

Immunoprecipitates were prepared from ~2×10^6^ GBM1 or HeLa cell pellets using either anti-TACC3 IgG1 ((D-2): sc-48368, Santa Cruz Biotechnology; dilution: 1:2000) or normal mouse control immunoglobulin (sc-3877, Santa Cruz Biotechnology; dilution: 1:2000). The protein precipitates were separated by SDS-PAGE followed by Coomassie staining. Distinct protein bands present solely in the ‘TACC3’ lane were excised and subject to trypsin digestion followed by liquid chromatography tandem mass spectrometry and protein identification as previously described (41). Protein-protein interaction networks were built via STRING v11 (23) based on text mining, experiments, databases, co-expression, neighbourhood, gene fusion, and co-occurrence with a confidence score = 0.4. Plots were generated with Cytoscape v3.8 (42).

### Gene expression profiling

#### Microarray

gene expression profiling was carried out using Human OneArray Plus (Phalanx Biotech Group). Target preparation was performed using an Eberwine-based amplification method with Amino Allyl MessageAmp II aRNA Amplification Kit (Ambion, AM1753) to generate amino-allyl antisense RNA (aa-aRNA). Labeled aRNA coupled with NHS-CyDye was prepared and purified prior to hybridization. Purified coupled aRNA was quantified using NanoDrop ND-1000; pass criteria for CyDye incorporation efficiency at > 15 dye molecular/1000nt. After microarray readout, raw data files were loaded into Rosetta Resolver® System (Rosetta Biosoftware) for profile error model calculation. The repeated probes within one chip were averaged resulting in a set of 28264 probes, and signal intensities were normalized using median scaling performed on data set (without flagged/control data).

#### Principal component (PCA) and single sample gene set enrichment analysis (ssGSEA)

RNAi off-target effects were alleviated by validating shRNA phenotypic redundancy across the probe set, filtering out all anticorrelated signals detected between TACC3 Kd1 and Kd2 (as determined by variance calculations across the cohort). Strictest cutoff criteria (no fold change limitation between anticorrelated probes) resulted in 3583 TACC3-dependent genes (Additional file S2: Correlated TACC3 Kd1 and Kd2 probes), that were used in the PCA analysis. PCA was performed within R (version 3.4.1) using the *prcomp* function of the *stats* package (version 3.4.1). Graphical rendering of the PCA (PC1-3) was performed using the pca3d package (version 0.1). Subsequently, ssGSEA was performed on a total of 231 pathways (including KEGG pathways and *TACC3*-containing pathways obtained from MSigDB (43) using the GSVA package (version 1.24.1), and selecting for geneset sizes between 10-500 genes. Subsequently, enrichment scores were computed for 142 pathways, for each of the two TACC3 knockdowns in relation to their respective (non-targeting) control per analyzed cell cycle stage (early G1, G1, S, G2/M; Table S2).

### Kohonen Self-Organizing feature map (SOM)

SOMs were generated using the Kohonen package version 3.0.10) under R (version 4.0.0) (44). Input features for the PATH_SOM constituted ssGSEA pathway enrichment scores across cell cycle stages. Phenotypically-correlated TACC3 Kd1 and Kd2 probes were used for the Gene_SOM. Data for each SOM was plotted with default parameters, and mapped using 2000 iterations (rlen=2000) against a 20-node rectangular map grid (xdim=4, ydim=5). Clustering was performed on SOM nodes; 10 clusters were estimated as the most suitable, using k-means clustering and examining “within cluster sum of squares”. Nodes were color-coded by cluster membership, and weight vector views (represented as inner pie charts per node) were used to indicate topological features of the cell cycle stages (early G1, G1, S, G2/M) and TACC3 knockdowns (Kd1 and Kd2).

### Survival and statistical analyses

Kaplan Meier survival plots with the respective hazard ratios and p-values for a given gene or gene signature were generated using GEPIA1 (source: http://gepia2.cancer-pku.cn/#survival) and GEPIA2 (source: http://gepia2.cancer-pku.cn/#survival) (28,35). ‘Overall’ and ‘Disease Free Survival’ (herein termed progression-free) functions were used with a median group cutoff and 95% confidence interval. The survival map using multiple datasets was generated using GEPIA2, with hazard ratio<0.05 and a median group cutoff. Default settings were used for Cutoff-high and cutoff-low percentages (50% each) for both survival plots and map. The Student’s t test (two-tailed, equal variance, n ≥ 3 independent experimental repeats) or Mann-Whitney U test (one-tailed) were used as stated in the caption of Fig. 1.

## Supplementary files

Supplementary file 1: Supplementary tables and figures (PDF)

Supplementary file 2: Correlated TACC3 Kd1 and Kd2 probes (CSV)

Supplementary file 3: ssGSEA results (CSV)

## Acknowledgements

We thank M. Lorger and P. Ceppi for proofreading the manuscript.

## Author’s contribution

HB, ESP, BKI, AZ, RKM, DMG, and HW designed and performed experiments, and analysed data. HW drafted the manuscript and HB, ESP, BKI, AZ, RKM, DMG, and HW edited the manuscript. ESP, RKM, DMG, and HW conceptualized the study. RKM, DMG, and HW supervised the study. All authors read and approved the final manuscript.

