## Supplementary file 1 for "The cell cycle state defines TACC3 as a regulator gene in glioblastoma"

### Supplementary tables and figures

### Supplementary tables

| Source | Weblink | Reference |
|---|---|---|
| THE HUMAN PROTEIN ATLAS | https://www.proteinatlas.org/ | (22) |
| Cytoscape | https://cytoscape.org/ | (42) |
| STRING v11 | https://string-db.org/ | (23) |
| MSig Database | https://www.gsea-msigdb.org/gsea/msigdb | (43) |
| KEGG Database | https://www.genome.jp/kegg/kegg1.html | (45) |
| GEPIA1 & GEPIA2 (Gene Expression Profiling Interactive Analysis) | http://gepia.cancer-pku.cn/ http://gepia2.cancer-pku.cn/ | (28)(35) |

**Table S1.** Use of publicly available data and open source software.

| CLUSTER | GENES |
|---|---|
| c25 | FGG,FXYD1,TMX3,ATP13A5,SHOX,CD33,CCKAR,CCNI2,PARVG,DNAJC15,MLIP,FAM110B,HEYL,LAMP3,CLLU1OS,IGSF1,LCE1E,FAM20A,OPTC,TMEM238,CDH5,PLAC8,ARFGEF3,FGL2,LRRC25,PGR,KCNMB4,ASPRV1,TLR7,APOBEC3H,DHRS9,NCOA2,CRYGD,CXCL12,CEACAM21,GAB1,CBX2,SMYD1,FGF17,ZBP1,DCN,SLC16A12,AIM2,SIGLEC11,ACKR3,C9orf139,CDHR3,IGSF21,RIMS3,H2BFWT,LOC100506801,NLRP7 |
| c32 | ALPL,RNF133,HIST3H2BB,GLYAT,ADAMTSL1,FGG,FXYD1,TMX3,ATP13A5,SHOX,CD33,CCKAR,CCNI2,PARVG,DNAJC15,MLIP,FAM110B,HEYL,LAMP3,CLLU1OS,IGSF1,LCE1E,FAM20A,OPTC,TMEM238,CDH5,PLAC8,ARFGEF3,FGL2,LRRC25,PGR,KCNMB4,ASPRV1,TLR7,APOBEC3H,DHRS9,NCOA2,CRYGD,CXCL12,CEACAM21,GAB1,CBX2,SMYD1,FGF17,ZBP1,DCN,SLC16A12,AIM2,SIGLEC11,ACKR3,C9orf139,CDHR3,IGSF21,RIMS3,H2BFWT,LOC100506801,NLRP7 |
| c31 | TCAF2,ANKRD39,P2RY2,TRPV3,TNFSF11,GPR143,ATP8B5P,CSN1S1,SLC37A2,NOS1,FBXO48,NGB,HBB,EVI5,SLC34A2,ADGRE2,CARD11,MYL4,FLT4,GUCA2A,COL17A1,NFE4,NR4A3,TRPC1,BAIAP2L2,DPEP3,FGD4,CMTM1,TCL1A,DAAM1,PCDH15,ADAT2,LOC643355,SERPINA11 |
| c21 | HOTAIR,PTPN22,HSD3B7,CYP11A1,CYP1B1,GDF2,SENP6,S100A12,PTPRM,COL2A1,MAPT-IT1,KCNAB3,POGZ,N4BP3,WFDC1,ERVFRD-1,FLT3,TPCN2,RARG,UNC13D,SCIMP,PLA2G2F,KRT3,RALGPS2,DQX1,LSAMP,COMMD6,KANK4,MRTO4,INHBC,SST,CLCNKA,LOC100130502,SLC30A8 |
| c40 | TMEM216,UPP1,NDST4,PLA2G5,HIPK3,HAAO,ARL8A,CLCN2,MFSD3,C20orf202,CYP1A2,EDNRA,ALDH7A1,ZPLD1,BRINP3,SAG,KANSL3,IL12A,MIEN1,MAP4K4,PSMA1 |
| c26 | HIST3H2A,P4HA2,FNDC3B,LILRA6,TXNIP,CCL2,C7orf31,FAM171B,ALOX5AP,ABCC2,NMUR1,SLA,CLNK,ABI3,C1orf115,CCDC136,AHSP,CXCL2,FAM179A,CSMD2,EPB41,TAS2R13,PRRG1,CDC20B,HAVCR1,NHS,CFAP44,SNRPN,CFTR |

**Table S2.** The 238 genes revealed by Gene_SOM clustering (comma delimited lists).

| **Symbol** | **Gene name** | **Logrank p-values** |
|---|---|---|
| HIST3H2A | Histone cluster 3 H2A | 0.00073 |
| P4HA2 | Prolyl 4-hydroxylase subunit alpha 2 | 0.0011 |
| TCAF2 | TRPM8 channel associated factor 2 | 0.0029 |
| HOTAIR | HOX transcript antisense RNA | 0.0063 |

| | | |
|---|---|---|
| GCKR | Glucokinase regulator | 0.0076 |
| ANKRD39 | Ankyrin repeat domain 39 | 0.01 |
| ALPL | Alkaline phosphatase, Biomineralization Associated | 0.011 |
| LOX | Lysyl oxidase | 0.013 |
| RNF133 | Ring finger protein 133 | 0.016 |
| SLC39A14 | solute carrier family 39 member 14 | 0.016 |
| PTPN22 | Protein tyrosine phosphatase non-receptor type 22 | 0.018 |
| HSD3B7 | Hydroxy-delta-5-steroid dehydrogenase, 3 beta- and steroid delta-isomerase 7 | 0.019 |
| FNDC3B | Fibronectin type III domain containing 3B | 0.019 |
| CSF3R | Colony stimulating factor 3 receptor | 0.025 |
| LILRA6 | Leukocyte immunoglobulin like receptor A6 | 0.026 |
| HIST3H2BB | Histone cluster 3 H2B family member b | 0.03 |
| P2RY2 | Purinergic receptor P2Y2 | 0.034 |
| GLYAT | Glycine-N-acyltransferase | 0.034 |
| ADAMTSL1 | ADAMTS like 1 | 0.035 |
| FGG | Fibrinogen gamma chain | 0.043 |
| TMEM216 | Transmembrane protein 216 | 0.047 |
| TRPV3 | Transient receptor potential cation channel subfamily V member 3 | 0.049 |

**Table S3.** All genes indicated by Gene_SOM whose elevated expression negatively correlates with overall survival in GBM with Logrank p-values < 0.05.

| Tumor number | Study abbreviation | Study name |
| --- | --- | --- |
| 1 | ACC | Adrenocortical carcinoma |
| 2 | BLCA | Bladder Urothelial Carcinoma |
| 3 | BRCA | Breast invasive carcinoma |
| 4 | CESC | Cervical squamous cell carcinoma and endocervical adenocarcinoma |
| 5 | CHOL | Cholangiocarcinoma |
| 6 | COAD | Colon adenocarcinoma |
| 7 | DLBC | Lymphoid Neoplasm Diffuse Large B-cell Lymphoma |
| 8 | ESCA | Esophageal carcinoma |
| 9 | GBM | Glioblastoma multiforme |
| 10 | HNSC | Head and Neck squamous cell carcinoma |
| 11 | KICH | Kidney Chromophobe |
| 12 | KIRC | Kidney renal clear cell carcinoma |
| 13 | KIRP | Kidney renal papillary cell carcinoma |
| 14 | LAML | Acute Myeloid Leukemia |
| 15 | LGG | Brain Lower Grade Glioma |
| 16 | LIHC | Liver hepatocellular carcinoma |

| | | |
|---|---|---|
| 17 | LUAD | Lung adenocarcinoma |
| 18 | LUSC | Lung squamous cell carcinoma |
| 19 | MESO | Mesothelioma |
| 20 | OV | Ovarian serous cystadenocarcinoma |
| 21 | PAAD | Pancreatic adenocarcinoma |
| 22 | PCPG | Pheochromocytoma and Paraganglioma |
| 23 | PRAD | Prostate adenocarcinoma |
| 24 | READ | Rectum adenocarcinoma |
| 25 | SARC | Sarcoma |
| 26 | SKCM | Skin Cutaneous Melanoma |
| 27 | STAD | Stomach adenocarcinoma |
| 28 | TGCT | Testicular Germ Cell Tumors |
| 29 | THCA | Thyroid carcinoma |
| 30 | THYM | Thymoma |
| 31 | UCEC | Uterine Corpus Endometrial Carcinoma |
| 32 | UCS | Uterine Carcinosarcoma |
| 33 | UVM | Uveal Melanoma |

**Table S4.** List of TCGA datasets analyzed.

| | |
|---|---|
| KEGG_GLIOMA | AKT1,AKT2,AKT3,ARAF,BRAF,CALM1,CALM2,CALM3,CALML3,CALML5,CALML6,CAMK2A,CAMK2B,CAMK2D,CAMK2G,CCND1,CDK4,CDK6,CDKN1A,CDKN2A,E2F1,E2F2,E2F3,EGF,EGFR,GRB2,HRAS,IGF1,IGF1R,KRAS,MAP2K1,MAP2K2,MAPK1,MAPK3,MDM2,MTOR,NRAS,PDGFA,PDGFB,PDGFRA,PDGFRB,PIK3CA,PIK3CB,PIK3CD,PIK3CG,PIK3R1,PIK3R2,PIK3R3,PIK3R5,PLCG1,PLCG2,PRKCA,PRKCB,PRKCG,PTEN,RAF1,RB1,SHC1,SHC2,SHC3,SHC4,SOS1,SOS2,TGFA,TP53 |
| PID_AURORA_A_PATHWAY | AJUBA,AKT1,ARHGEF7,AURKA,AURKAIP1,AURKB,BIRC5,BRCA1,CDC25B,CENPA,CKAP5,CPEB1,DLGAP5,FZR1,GADD45A,GIT1,GSK3B,MDM2,NDEL1,NFKBIA,OAZ1,PAK1,PPP2R5D,PRKACA,RAN,RASA1,TACC1,TACC3,TDRD7,TP53,TPX2 |
| FISCHER_G1_S_CELL_CYCLE | ABCA5,ABHD10,ABHD4,ACYP1,ADAMTS1,ADCK2,ADCY6,AHI1,AKAP5,AMIGO2,ANK3,ANKRA2,AP3M2,ARGLU1,ARID5B,ARL6IP6,ASPH,ATAD2,BARD1,BBX,BLM,BRCA1,BRIP1,C1GALT1,CASP2,CASP8AP2,CCN2,CCND3,CCNE1,CCNE2,CDC25A,CDC6,CDC7,CDCA7,CDH10,CDH24,CDT1,CENPQ,CHAF1A,CHAF1B,CHEK1,CLSPN,CNNM3,COL4A4,CTPS1,DCLRE1A,DCLRE1B,DLGAP1,DNAJC3,DNAJC6,DSCC1,DTL,DUSP10,E2F1,E2F2,E2F7,E2F8,EAF2,EXO1,EZH2,FAM111B,FANCE,FANCG,FEN1,FOSB,FST,GCLC,GINS2,GK,GLCCI1,GMNN,HELLS,HSPB8,ICMT,IGF1R,INSIG2,INSR,IVNS1ABP,JUND,KLF5,KLRK1,LCMT2,LIPH,MAGEL2,MAN1A2,MAP2K6,MASTL,MATN1,MBNL2,MCM10,MCM2,MCM3,MCM5,MCM6,MDM1,MITF,MNS1,MNT,MNX1,MRC2,MSH2,MSH5,MSH6,MYB,NASP,NCOA7,NEDD9,NPAT,NR4A3,NUDT7,OGT,OPCML,OSBPL6,OSGIN2,PANK2,PAQR4,PASK,PCNA,PEX11B,PEX13,PLCXD1,PLSCR4,POLA2,POLD3,POLE,POLE2,PRIM1,PRKD1,PTGS2,RABIF,RAD51,RBBP8,RECQL4,REEP1,RFC2,RFC4,RFXAP,RGS7,RIMKLB,RMI1,RNPC3,RPA2,RRM2,SLBP,SLC25A27,SLC38A2,SMPD1,SP1,SPIN3,SSBP2,SVIP,TAF15,TCEAL9,TCF19,TENM3,TIFA,TIPIN,TLR3,TLR6,TMCC1,TMEM243,TNS2,TOPBP1,TP53INP1,TREX1,TRIM45,TTLL7,TXNRD1,UBR7,UNG,USP1,USP18,USP37,USP53,WDHD1,WDR76,YEATS4,ZBTB14,ZMYND19,ZNF367,ZRANB2 |
| JOHANSSON_GLIOMAGENESIS_BY_PDGFB_UP | B2M,C1orf159,CCND1,CCND2,CD63,CD9,CDC20,CDC42SE1,CDK1,CDK4,CHST11,DBI,DYNLT1,EEF1A1,EPN2,FN1,FOS,FRMD8,FTL,HLADRB5,HMGB2,HMOX1,HNRNPA1,HSPA14,KLF6,LGALS1,MARCKS,MARCKSL1,MCAM,MCM2,MCM3,MKI67,NIN,OST4,PBK,PCNA,PDGFRA,PLK1,PPFIBP1,PPP1R14B,PPP1R18,REV3L,RNF213,RPLP0,RPS27L,RRM2,SDC3,SERPINE2,SPP1,STAT3,SULF2,TAGLN2,TMEM176B,TOP2A,TPM1,TPM4,YBX1 |
| TANG_et_al, based on (21) | ABCA1,AEBP1,ALOX5AP,CD14,CD163,CD44,CFI,CHI3L2,CLIC1,COL1A1,COL1A2,CXCR4,ECM2,FCER1G,FNDC3B,GPNMB,RING6,HMOX1,IFI44,IGFBP2,IGFBP3,LY96,MMP |

| | |
|---|---|
| | 2,MTHFD2,MYD88,NMI,PLSCR1,PTX3,PXDN,PYGL,RBBP8,SERPINE1,SOD2,SRPX,FAM46A,TGFB1,TIMP1,VSIG4 |
| HALLMARK_G2M_CHECKPOINT | ABL1,AMD1,ARID4A,ATF5,ATRX,AURKA,AURKB,BARD1,BCL3,BIRC5,BRCA2,BUB1,BUB3,CASP8AP2,CBX1,CCNA2,CCNB2,CCND1,CCNF,CCNT1,CDC20,CDC25A,CDC25B,CDC27,CDC45,CDC6,CDC7,CDK1,CDK4,CDKN1B,CDKN2C,CDKN3,CENPA,CENPE,CENPF,CHAF1A,CHEK1,CHMP1A,CKS1B,CKS2,CTCF,CUL1,CUL3,CUL4A,CUL5,DBF4,DDX39A,DKC1,DMD,DR1,DTYMK,E2F1,E2F2,E2F3,E2F4,EFNA5,EGF,ESPL1,EWSR1,EXO1,EZH2,FANCC,FBXO5,FOXN3,G3BP1,GINS2,GSPT1,H2AX,H2AFZ,H2AFV,H2BFT,HIF1A,HIRA,HMGA1,HMGB3,HMGN2,HMMR,HNRNPD,HNRNPU,HOXC10,HSPA8,HUS1,ILF3,INCENP,ARM2,KATNA1,KIF11,KIF15,KIF20B,KIF22,KIF23,KIF2C,KIF4A,KIF5B,KMT5A,KNL1,KPNA2,KPNB1,LBR,LIG3,LMNB1,MAD2L1,MAPK14,MARCKS,MCM2,MCM3,MCM5,MCM6,MEIS1,MEIS2,MKI67,MNAT1,MT2A,MTF2,MYBL2,MYC,NASP,NCL,NDC80,NEK2,NOLC1,NOTCH2,NSD2,NUMA1,NUP50,NUP98,NUSAP1,ODC1,ODF2,ORC5,ORC6,PAFAH1B1,PBK,PDS5B,PLK1,PLK4,PML,POLA2,POLE,POLQ,PRC1,PRIM2,PRMT5,PRPF4B,PTTG1,PTTG3P,PURA,RACGAP1,RAD21,RAD23B,RAD54L,RASAL2,RBL1,RBM14,RPA2,RPS6KA5,SAP30,SFPQ,SLC12A2,SLC38A1,SLC7A1,SLC7A5,SMAD3,SMARCC1,SMC1A,SMC2,SMC4,SNRPD1,SQLE,SRSF1,SRSF10,SRSF2,SS18,STAG1,STIL,STMN1,SUV39H1,SYNCRIP,TACC3,PAPD7,TFDP1,TGFB1,TLE3,TMPO,TNPO2,TOP1,TOP2A,TPX2,TRA2B,TRAIP,TROAP,TTK,UBE2C,UBE2S,UCK2,UPF1,WRN,XPO1,YTHDC1 |
| KEGG_CELL_CYCLE | ABL1,ANAPC1,ANAPC10,ANAPC11,ANAPC13,ANAPC2,ANAPC4,ANAPC5,ANAPC7,ATM,ATR,BUB1,BUB1B,BUB3,CCNA1,CCNA2,CCNB1,CCNB2,CCNB3,CCND1,CCND2,CCND3,CCNE1,CCNE2,CCNH,CDC14A,CDC14B,CDC16,CDC20,CDC23,CDC25A,CDC25B,CDC25C,CDC26,CDC27,CDC45,CDC6,CDC7,CDK1,CDK2,CDK4,CDK6,CDK7,CDKN1A,CDKN1B,CDKN1C,CDKN2A,CDKN2B,CDKN2C,CDKN2D,CHEK1,CHEK2,CREBBP,CUL1,DBF4,E2F1,E2F2,E2F3,E2F4,E2F5,EP300,ESPL1,FZR1,GADD45A,GADD45B,GADD45G,GSK3B,HDAC1,HDAC2,MAD1L1,MAD2L1,MAD2L2,MCM2,MCM3,MCM4,MCM5,MCM6,MCM7,MDM2,MYC,ORC1,ORC2,ORC3,ORC4,ORC5,ORC6,PCNA,PKMYT1,PLK1,PRKDC,PTTG1,PTTG2,RAD21,RB1,RBL1,RBL2,RBX1,SFN,SKP1,SKP2,SMAD2,SMAD3,SMAD4,SMC1A,SMC1B,SMC3,STAG1,STAG2,TFDP1,TFDP2,TGFB1,TGFB2,TGFB3,TP53,TTK,WEE1,WEE2,YWHAB,YWHAE,YWHAG,YWHAH,YWHAQ,YWHAZ,ZBTB17 |
| JOHANSSON_ BRAIN_CANCER_ EARLY_VS_LATE | ACOT7,ALDOC,ATP1B1,ATP2A2,CALM1,CALY,CAMK2G,CHN1,CLSTN1,CLU,CRYAB,CSDC2,CST3,CTSD,DCN,GHITM,GJA1,GNAS,GRINA,HDAC11,IGFBP4,IGFBP5,ITM2B,LRP1,MAT2B,MMD2,NCDN,NDN,NDRG1,NDRG2,NSF,PFKM,PLA2G7,PLEKHB2,PYGB,QDPR,SELENOP,SPARC,SPARCL1,TEF |

| | |
|---|---|
| KIM_et_al, based on (22) | DEPTOR,RPRM,NET1,WAC,RNF178,REPS2,ZNF609,KLF13,CXCL8,ADM,PDPN,IGFBP2,MDK,TIMP1,EFEMP2,ACOX2,TAGLN2,SLC43A3,LGALS8,DYNLT3,KIAA0323,TFRC,FBXO17,SLC35G2,PINLYP,MT1E,DCTD,HOMER1,FAM3C,CASP3,NSUN5,PDLIM3,MT1M |
| TACC3_GSOM | P4HA2,FNDC3B,LILRA6,TMEM216,TCAF2,ANKRD39,P2RY2,TRPV3,HOTAIR,PTPN22,HSD3B7,ALPL,RNF133,HIST3H2BB,GLYAT,ADAMTSL1,FGG,GCKR,LOX,SLC39A14,CSF3R |

**Table S5.** Signatures used for survival analysis (comma delimited lists).

**Supplementary figures**

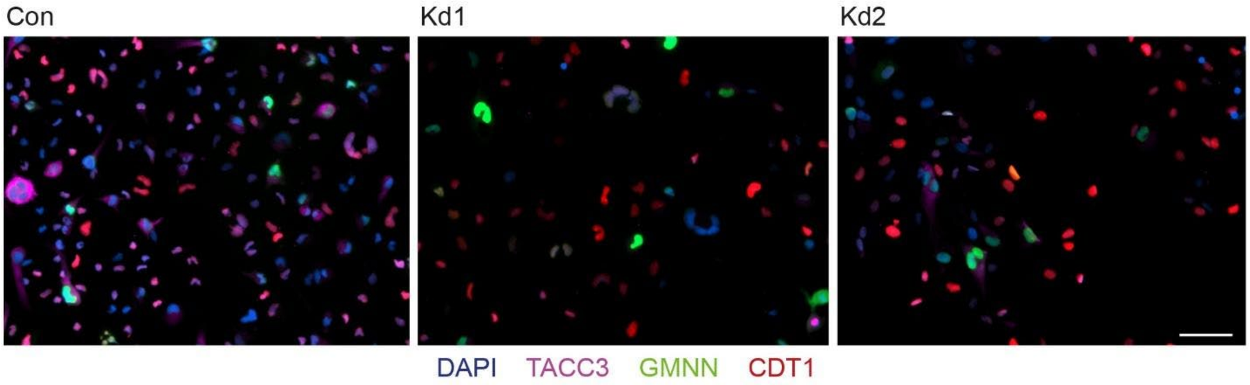


**Figure S1.** Images of 1_FUCCI control and TACC3 knockdown (Kd1, Kd2) cells. Cells were stained with DAPI. Scale bar, 50 μm.

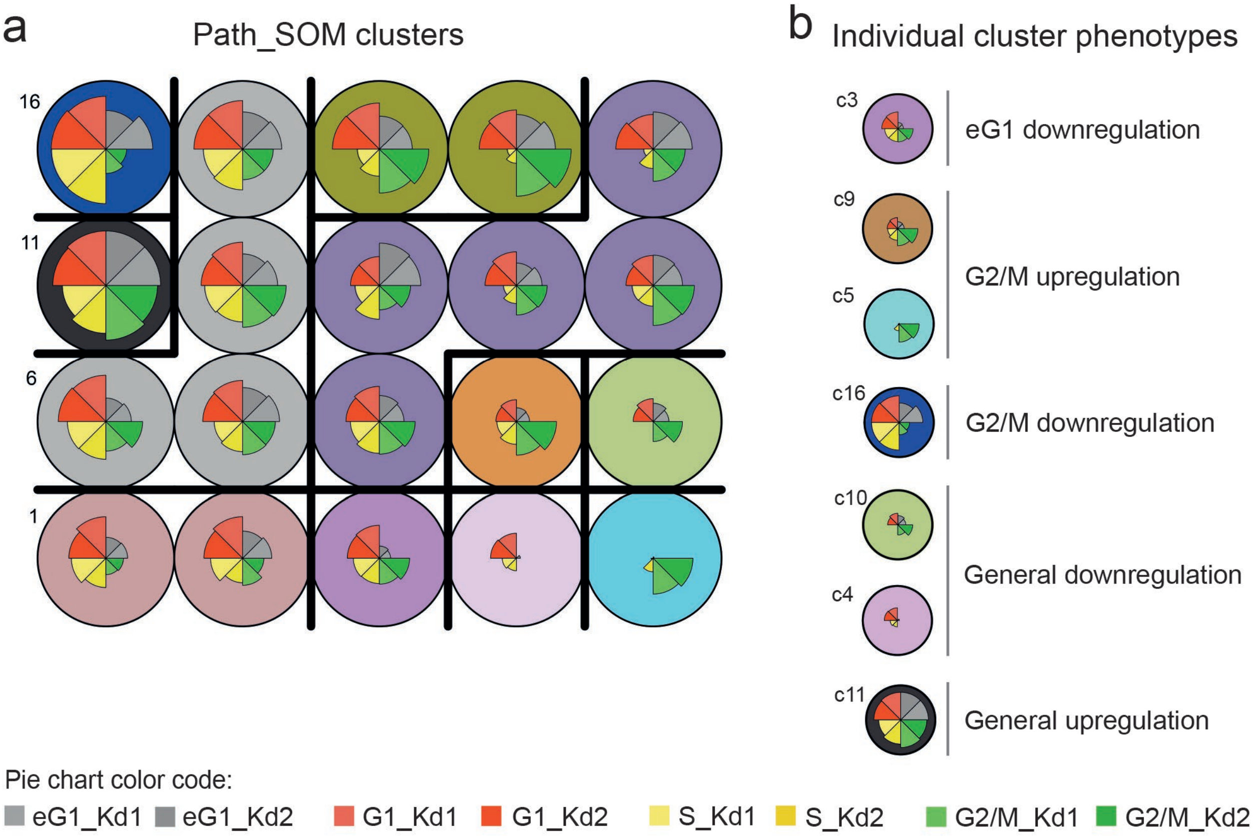


**Figure S2.** Results of Path_SOM. **A)** Data visualization of the SOM map grid containing 20 nodes (numbered 1-20), based on pathway enrichment patterns across cell cycle stages (indicated by inner pie charts per node). Nodes are colored by cluster membership, and identified clusters (n=10) are delineated by black lines. **B)** Categorization of individual PATH_SOM clusters according to the indicated cell cycle phenotype (pie charts).

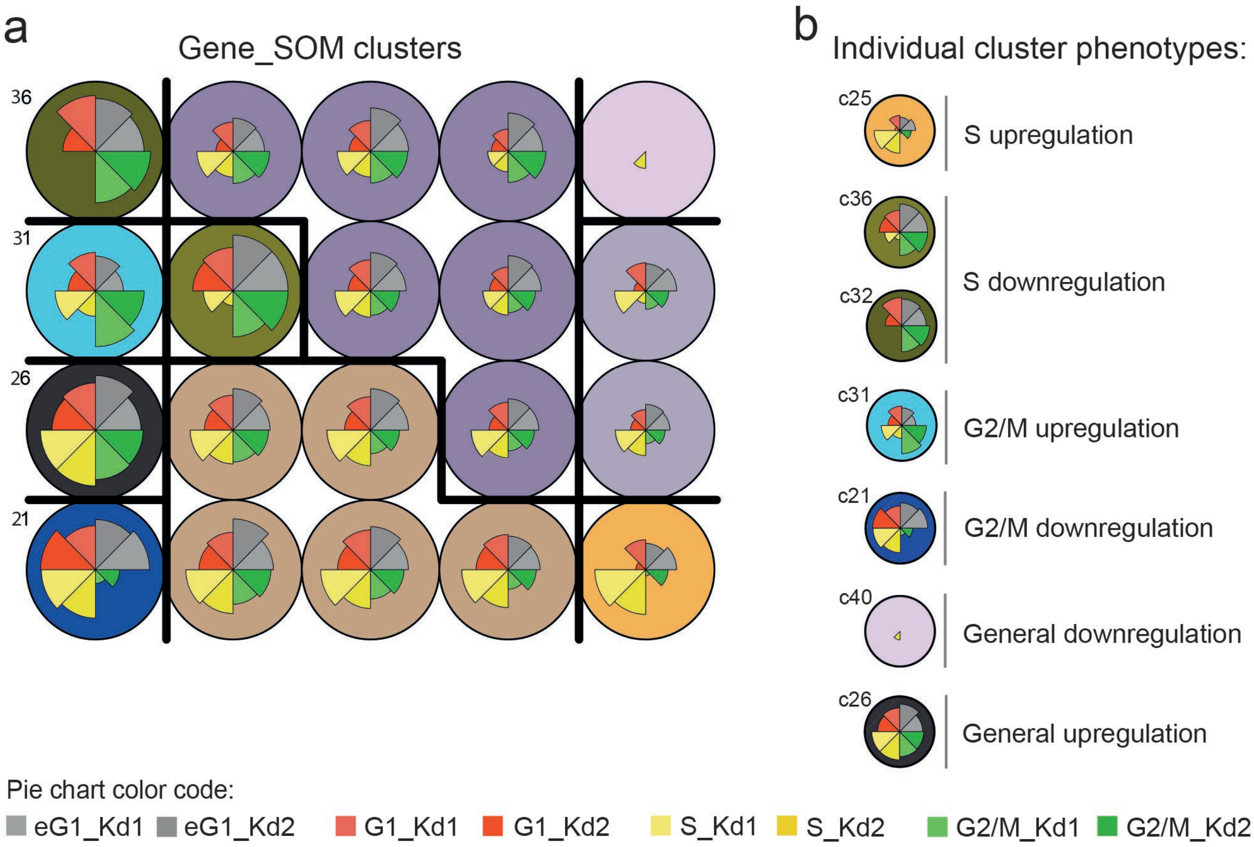


**Figure S3.** Results of Gene_SOM. **A)** Data visualization by 20 SOM nodes (numbered 21-40) and 10 clusters (indicated by black lines). **B)** Categorization of individual PATH_SOM clusters according to the indicated cell cycle phenotype (pie charts).
